# Cooperative dynamics of DNA grafted magnetic nanoparticles optimize magnetic biosensing and coupling to DNA origami

**DOI:** 10.1101/2023.04.11.536349

**Authors:** Aidin Lak, Yihao Wang, Pauline J. Kolbeck, Christoph Pauer, Mohammad Suman Chowdhury, Marco Cassani, Frank Ludwig, Thilo Viereck, Florian Selbach, Philip Tinnefeld, Meinhard Schilling, Tim Liedl, Joe Tavacoli, Jan Lipfert

## Abstract

Magnetic nanoparticles (MNPs) enable unique capabilities for biosensing and actuation via coupling to DNA origami, yet how DNA grafting density affects their dynamics and accessibility remains poorly understood. Here, we demonstrate functionalization of MNPs with single-stranded DNA (ssDNA) via click chemistry conjugation with tunable grafting density. Several complementary methods show that particle translational and rotational dynamics exhibit a sigmoidal dependence on ssDNA grafting density. At low densities ssDNA strands are coiled and cause small changes to particle dynamics, while at high densities they form polymer brushes that cooperatively change particle dynamics. Intermediate ssDNA densities show the highest magnetic biosensing sensitivity for detection of target nucleic acids. Finally, we demonstrate that MNPs with high grafting densities are required to efficiently couple them to DNA origami. These results together establish ssDNA grafting density as a critical parameter for functionalization of MNPs for use in a broad range of applications.

## INTRODUCTION

DNA grafted inorganic nanoparticles are versatile building blocks for superstructures of selected geometries and functionalities as well as responsive nanomaterials for biosensing and biocomputing.^1–5^ Programmed assemblies of gold (Au) nanoparticles (NPs) using Watson-Crick base pairing with complementary single-stranded DNA (ssDNA) strands were introduced two decades ago.^1, 2^ DNA labelled NPs and nanostructures have been used as molecular nanothermometers,^6^ *in vitro* biosensors,^7^ nanoparticle beacons,^8^ and as an intracellular sensing platform.^9^ NP-DNA conjugates offer novel possibilities for further modifications through competitive hybridization,^10^ DNA strand displacement,^7^ and ligation.^11^ The functionalization of Au NPs with ssDNA can be achieved through direct Au-S covalent bonds, making it straightforward and requiring only minimal surface modification steps. Consequently, Au NP-DNA conjugates have been combined with DNA origami nanotechnology to develop novel tools and applications, for example chiral plasmonic nanostructures, plasmonic rulers, and nanoantennas for biosensing.^12–18^

Colloidal magnetic nanoparticles (MNPs) transduce magnetic energy to heat under alternating magnetic fields, a feature that is harnessed in magnetic hyperthermia-triggered drug delivery and magnetic actuation of cellular processes.^19–21^ Combining the potential for local heating by external fields with the well-defined melting temperature of double-stranded DNA (dsDNA) has led to molecular magnetic nanothermometers.^6^ Further, MNPs change their Brownian magnetic relaxation dynamics upon binding to targeting molecules or in response to immediate environmental cues such as viscosity and temperature.^22^ Building upon this feature, magnetic biosensing has become a versatile platform for rapid, mix-and-measure, and quantitative detection of bacteria,^23^ proteins,^24, 25^ and DNA/RNA.^26–28^ Despite offering all these unique features in a single entity, DNA-based magnetic nanoparticles and nanostructures are still poorly studied and explored. This knowledge gap mainly stems from the multistep pre-modification and surface chemistries that are needed for successful DNA labelling of MNPs. In contrast to Au NPs, an intermediate encapsulation layer with a well-defined chemistry, functionality, and organization is required to bind ssDNA to MNPs.^29^ Surface silanization is commonly used to pre-modify the particle surface.^30^ Electrostatic interactions between charged nanoparticles and dsDNA macromolecules have been successfully used to attached dsDNA to MNPs, yet only after a multistep surface silanization.^31^ However, the accessibility and targeting ability of ssDNA bound via unspecific charge interactions is rather poor due to the binding of DNA to multiple positive head groups, causing them to flatten on the particle surface. Amine-modified ssDNA have been covalently conjugated to MNPs,^32, 33^ again after surface silanization.^34, 35^ Encapsulation of iron oxide NPs in an amphiphilic polymer that offers copper-free click chemistry conjugation has enabled their assembly into superlattices^36^ and binding to DNA origami structures.^37^ Yet our understanding about how the DNA grafting density changes the particle dynamics, determines the accessibility of ssDNA to complementary sequences, and dictates the optimal application of MNPs is still limited.

Here, we demonstrate clickable magnetic nanoparticles (CMPs) offering copper-free click chemistry conjugation as a highly versatile nanoplatform to tune the grafting density of ssDNA over a broad range. We show that magnetic relaxation, hydrodynamic, and electrophoretic dynamics of CMP-DNA conjugates change in a cooperative manner in response to ssDNA grafting, which can be described by the Hill equation. Whereas at low grafting density the ssDNA strands are coiled on CMPs and minimally influences the dynamics of CMPs, at high grafting density they form dense polymer brushes and cooperatively change the dynamics of CMPs. Exploiting our CMPs as nanomarkers for magnetic-based biosensing, we find that CMP-DNA conjugates at the ssDNA grafting density corresponding to the midpoint of the Hill equation reveal the largest magnetic signal change upon hybridization to the target sequence. We propose that the ssDNA strands are in a mixture of coiled and brushed states at the intermediate grafting density. This configuration leads to the largest change in magnetic relaxation dynamics of particles upon duplexing with the target nucleic acids. In addition, we demonstrate coupling of our CMP-DNA conjugates to six-(6 HB) and 24-helix-bundle (24 HB) DNA origami structures. We find that high ssDNA grafting density favors binding to DNA origami structures with high efficiency and site-specificity.

## RESULTS AND DISCUSSION

### Polymer encapsulated magnetic nanoparticles offering high colloidal stability and copper-free click chemistry conjugation

To investigate the effects of ssDNA grafting density on particle dynamics and their accessibility toward target DNA strands with magnetic techniques, we use MNPs relaxing via a Brownian relaxation mechanism. We, therefore, synthesized cubic-shape cobalt-doped iron oxide MNPs using a high temperature decomposition synthesis procedure (see Methods for details).^38^ The stoichiometry was determined to be Co_0.85_Fe_2.15_O_4_ by inductively-coupled plasma optical emission spectroscopy (ICP-OES) and the mean particle side length *L*_c_ to be 15.8 ± 1.2 nm (mean ± standard deviation) by transmission electron microscopy (TEM; Figure 1a, c).

**Figure 1.**
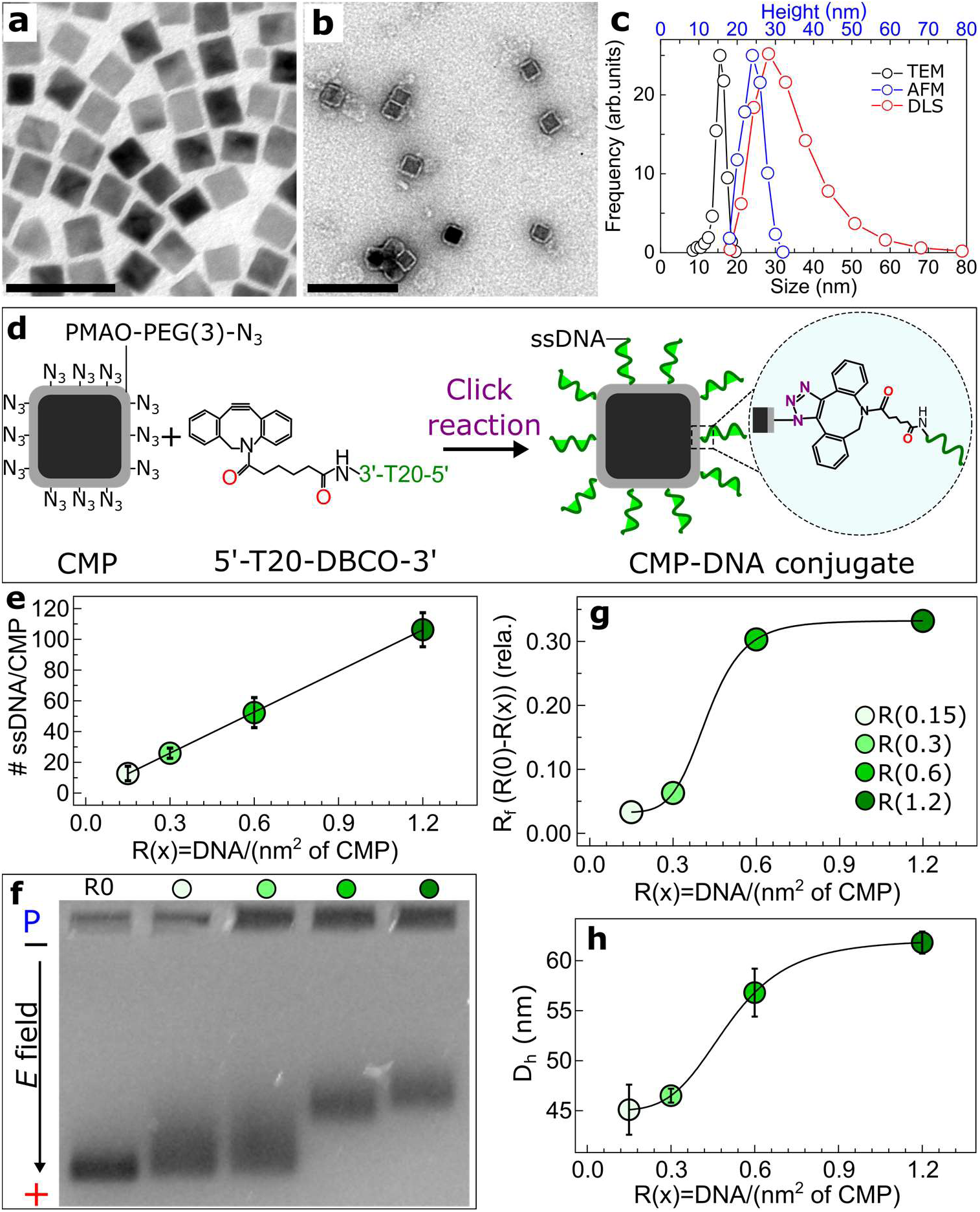
Functionalization and characterization of oleic acid coated MNPs, CMPs, and CMP-DNA conjugates. (a) Typical transmission electron microscopy (TEM) micrograph of oleic acid coated MNPs in chloroform. (b) Negative-stain (1% uranyl formate) TEM micrograph of PMAO-PEG(3)-azide coated MNPs (CMPs) before ssDNA labelling. (c) Side length *L*_c_ of oleic acid coated MNPs (from panel a), height *H* (AFM), and hydrodynamic diameter *D*_h_ histograms of CMPs after polymer coating. The height histogram was acquired from analysis of the AFM micrograph (Figure S1). The *D*_h_ distribution was recorded on Malvern-Zetasizer-Nano instrument on CMPs dispersed in Tris-EDTA (TE) buffer (5 mM Tris, 1 mM EDTA, 5 mM NaCl, pH 7.3). (d) Scheme of the strain-promoted azide-alkyne cycloaddition (SPAAC) copper-free click reaction between azide labelled CMPs and dibenzocyclooctyne (DBCO)-modified ssDNA oligomers. (e) The average number of ssDNA per CMP ± standard error of the mean (SEM) as estimated from the depletion assays *vs.* the nominal grafting density R(x). (f) Agarose gel electrophoresis shift assays performed on CMPs and CMP-DNA conjugates at different R(x) values on a 0.5% agarose gel at 90 mA, P stands for loading pockets. (g) The gel relative front (*R*_f_) of CMP-DNA conjugates relative to *R*_f_ of the R(0) sample with no ssDNA as a function of R(x). (h) The volume-weighted particle hydrodynamic size (*D*_h_) of CMP-DNA conjugates ± SEM as a function of R(x). Solid lines in panel f and g are fits of the Hill equation with fitted parameters *R*_half_ = 0.42 DNA/nm^2^ and *n* = 6.3 and *R*_half_ = 0.48 DNA/nm^2^ and *n* = 4.5, respectively. The scale bars are 50 and 100 nm in panels (a) and (b), respectively.

The MNPs were transferred to aqueous phase applying the “grafting to” polymer coating approach using poly(maleic anhydride-*alt*-1-octadecene) (PMAO) copolymers by adopting the procedure by Pellegrino et. al.^39^ To obtain singly-coated and “clickable” MNPs (CMPs), we first modified PMAO copolymers with polyethylene glycol(PEG)(3)-azide linkers and further optimized the original polymer coating recipe (see Methods for details). The conjugation of NH_2_-PEG(3)-azide linkers to PMAO was carried out through an anhydride ring opening reaction in dimethylformamide.^40^ During the polymer coating procedure and by gradual removal of chloroform, the C16 carbon chains of PMAO interact hydrophobically with the C18 chains of oleic acid and form a dense shell around MNPs. To visualize the polymeric shell and examine its uniformity and morphology, we performed negative-stain TEM imaging (Figure 1b). We observed that MNPs are uniformly coated with a ∼3 nm layer of the PMAO-PEG polymer. Moreover, most CMPs are singly enveloped inside the polymeric shell after the coating process. An interesting observation is that the PMAO-PEG copolymers shell around MNPs follows the cubic shape of particles (Figure 1b). Atomic force microscopy (AFM) images of CMPs deposited on mica substrate by adding Na^+^ to particle suspensions show that the CMPs are monodisperse and sparsely distributed on the substrate (Figure S1). From analysis of AFM images, we find that the particle height *H* increases from *L*_c_ = 15.8 ± 1.2 nm (from TEM) to *H* = 24.0 ± 2.6 nm (mean ± standard deviation (std)) after polymer coating (Figure 1c). Considering 1.8 nm and ∼1 nm for the length of a C18 chain of oleic acid and a PEG linker with 3 units, respectively,^41^ we expect an increase in size after the polymer coating by 2·1.8 nm + 2·1 nm = 5.6 nm, in good agreement with the experimentally observed difference between *H* and *L*_c_ of 8.2 ± 3.8 nm. Dynamic light scattering (DLS) reveals that the CMPs are colloidally stable with no detectable aggregates (Figure 1c). The number-weighted particle hydrodynamic diameter *D*_h_ is 32.3 ± 8.8 nm with a polydispersity index (PDI) of 0.1. The 10 nm difference between *D*_h_ and *H* is likely due to hydration and an electrical double layer that is formed around the charged CMPs in solution.

### The number of ssDNA per nanoparticle increases linearly with nominal grafting density

Our single-core CMPs offer an excellent nanoplatform to tune the grafting density of ssDNA to study its effect on the dynamic properties of the particles. We use the strain-promoted azide-alkyne cycloaddition (SPAAC) copper-free click chemistry to efficiently graft dibenzocyclooctyne (DBCO)-modified ssDNA strands of 20-mer thymine (T)_20_ on CMPs (Figure 1d). We designed our DNA labelling experiments using the grafting density parameter R(x), defined by the nominal number of ssDNA per nm^2^ of CMP, using the concentrations of CMPs and ssDNA in the reaction mixture and assuming the particle surface area to be 6.(^D^_h_/3)^2^ (see the discussion in Methods). Accordingly, we synthesized CMP-DNA conjugates at four different R(x) values of 0.15, 0.3, 0.6, and 1.2 ssDNA per nm^2^ of CMP. To determine the actual number of ssDNA per CMP at each nominal R(x), we performed depletion assays by measuring UV absorption spectra of supernatants of freshly DNA-labelled CMPs after centrifugation and particle pelleting. Three sequential supernatants after particle re-dispersion and centrifugation were measured. We converted the absorption at 260 nm to the number of ssDNA using the Beer-Lambert equation after correcting for background from particles (see Methods for protocols and equations). Remarkably, we found that the average number of ssDNA strands per CMP increases linearly with the nominal density R(x) (Figure 1e). Our data suggest that up to the highest nominal ssDNA density used, steric hindrance and electrostatic repulsion from the bound strands do not significantly inhibit further attachment of ssDNA to the CMPs. In the following, we consistently report the nominal density R(x) for ease of comparison, but note that R(x) can be directly scaled to the experimental density using the linear relationship in Figure 1e.

### Electrophoretic mobility and hydrodynamic size of CMP-DNA conjugates reveal a non-linear dependence on ssDNA grafting density

To characterize CMPs and CMP-DNA conjugates, we first performed agarose gel electrophoresis shift assays. All particles migrate towards the anode as a single band due to their overall negative charge (Figure 1f). We observe that the mobility of CMP-DNA conjugates in the gel decreases with increasing R(x). To parametrize the gel shift assay, we analyze the gel and extract relative front *R*_f_ values that indicate the migration of CMP-DNA conjugates relative to CMPs (without DNA functionalization). The *R*_f_ vs. R(x) is well described by a Hill equation with *R*_half_ = 0.42 DNA/nm^2^ and *n* = 6.3 (Figure 1g, the Hill equation is given in Methods). The particle hydrodynamic size *D*_h_ measured by DLS reveals a similar non-linear trend with R(x) that can be described by a Hill fit with *R*_half_ = 0.48 DNA/nm^2^ and *n* = 4.5 (Figure 1h). Our data reveal the presence of highly cooperative dynamics between the ssDNA strands on CMPs. Whereas the average number of ssDNA per CMP increases linearly (Figure 1e), the hydrodynamic and electrophoretic properties of CMP-DNA conjugates change in a non-linear and cooperative manner.

### Magnetic relaxation of CMPs changes cooperatively with grafting density

Next, we investigate whether the cooperative behavior of ssDNA on CMPs is reflected in the magnetic relaxation dynamics of particles. To monitor changes in magnetization dynamics of CMPs after DNA grafting, we performed complex magnetic ac-susceptibility (ACS) measurements on particle suspensions in Tris-EDTA (TE) buffer, which monitor magnetic relaxation dynamics of particles as a function of excitation frequency. The imaginary parts of the ACS spectra of CMPs and CMP-DNA conjugates show peaks that are typical of Brownian relaxation processes (Figure 2a).^42^ The shift in the Brownian peak position toward lower frequencies indicates a slower/retarded relaxation process, corresponding to a larger hydrodynamic volume, with increasing grafting density (Figure 2a,b). We find that the relaxation peak frequency *f*_p_ shifts as a function of R(x) in a non-linear manner (Figure 2c). Similar to *R*_f_ and *D*_h_, the dependence of the particle relaxation peak frequency *f*_p_ on R(x) is well described by a Hill equation, with *R*_half_ = 0.42 DNA/nm^2^ and *n* = 3.6, again suggesting cooperative changes in the magnetic relaxation dynamics of CMPs upon ssDNA grafting.

**Figure 2.**
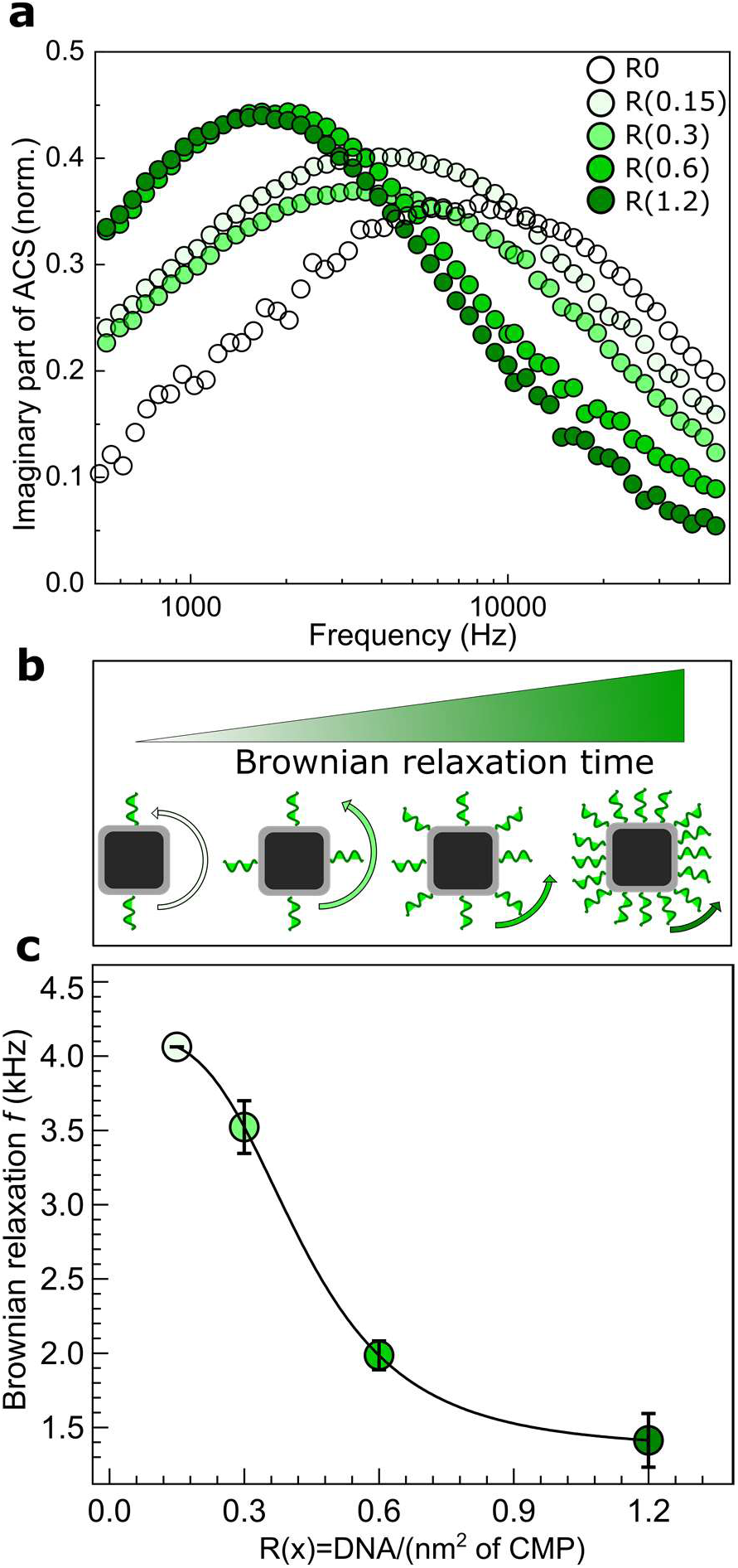
Characterization of magnetic relaxation of CMPs and CMP-DNA conjugates by complex magnetic ac-susceptibility (ACS) measurements. (a) Imaginary part of complex ac magnetic susceptibility spectra of the R(0) to R(1.2) samples, showing the shift in the characteristic Brownian magnetic relaxation peak of CMPs after ssDNA grafting. (b) Schematic of the nanoparticles at different grafting density, indicating how the particle Brownian relaxation frequency is reduced when more ssDNA are grafted to the CMPs. (c) Shift in the relaxation frequency *f*_p_ (mean ± SEM) after ssDNA grafting with respect to the R(0) particles having no ssDNA vs. R(x). The ACS measurements were performed at µ_0_*H* = 95 µT at 22 °C on 150 µl of particle suspensions in TE buffer. The black line is a fit of the Hill model, with parameters *R*_half_ = 0.42 DNA/nm^2^ and *n* = 3.6.

We hypothesize that ssDNA strands go through a transition from being coiled at low grafting density toward forming polymer brushes at high grafting density. Notably, we find a pronounced non-linear dependence on R(x) with similar *R*_half_ and *n* parameters from fitting the Hill model to the results of electrophoretic mobility, DLS, and magnetic relaxation measurements. The three analysis techniques rely on completely different working principles, but all three provide measurements of the particle size in solution.

### DNA grafting increases the colloidal stability of CMPs in the presence of Mg_2+_

To further characterize the grafted DNA shell and its response to environmental conditions, we added monovalent NaCl or divalent MgCl_2_ salt to the TE coupling buffer and probed our CMP-DNA conjugates. For detailed characterization, we used two complementary techniques: AFM imaging of nanoparticles deposited and dried on a mica substrate (Figure 3a-d), and DLS, which measures particles in solution. Due to the high negative charge density of DNA, both ionic strength and ion valency are expected to have significant effects on the DNA-coated nanoparticles.^43^

**Figure 3.**
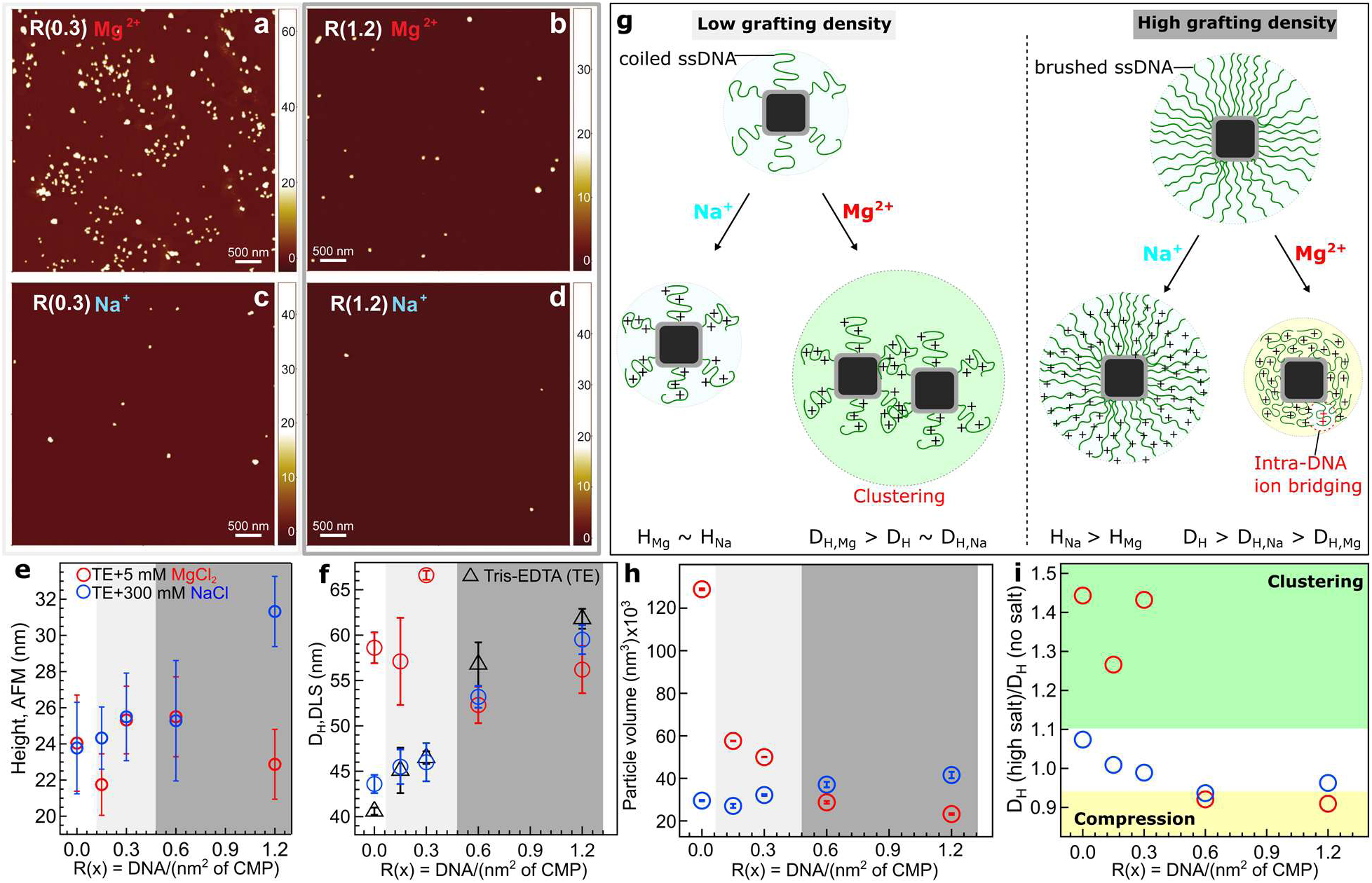
Characterization of CMP-DNA conjugates by atomic force microscopy and dynamic light scattering in the presence of monovalent or divalent salt. (a) and (c) AFM micrographs of low ssDNA grafted sample R(0.3) at 5 mM MgCl_2_ and 300 mM NaCl concentration, respectively. (b) and (d) AFM micrographs of high ssDNA grafted sample R(1.2) in the same ionic conditions. (e) Particle heights determined from AFM as a function of R(x). The data points are the mean ± std. Only single particles are selected for analysis. (f) Volume-weighted particle hydrodynamic size *D*_h_ (mean ± SEM) of CMP-DNA conjugates vs. R(x) measured by DLS in TE buffer only, with 5 mM MgCl_2_ or with 300 mM NaCl. (g) Schematic illustration of CMP-DNA conjugates at low and high ssDNA grafting densities in the presence of monovalent Na^+^ or divalent Mg^2+^ ions. (h) particle volume analysis (mean ± SEM) of the AFM images. Here, clusters are also included in the analysis. (i) Ratio of *D_h_* at high to low/no salt concentration vs. R(x), revealing the compression and clustering regimes in the presence of Mg^2+^ ions. The same color code is applied to panels (e), (f), (h), and (i). The low and high grafting density regimes are highlighted with light and dark gray colors in panels (e), (f), and (h).

Under high Na^+^ conditions, we find monodisperse and sparsely distributed CMPs on the mica substrate used for AFM imaging for all grafting densities (Figure 3c,d and Figure S1). The particle height *H* from the AFM images (Figure 3e and Figure S2) and the hydrodynamic diameter *D*_h_ from DLS show a consistent trend for the Na^+^ condition: the size of the particles increases with increasing grafting density, in line with the findings in low salt TE buffer (Figures 1h and 3f and Figure S3). As expected, the *D*_h_ values from DLS are consistently larger than the heights determined by AFM under dried conditions, in agreement with a view that the DNA brushes swell in solution and that the overall solvation layer extends significantly beyond the particles’ surface. This coincides also with the observation that DNA appears significantly more compact when imaged under dried conditions compared to in liquid imaging.^44^

For the Mg^2+^ condition, AFM imaging reveals the formation of particle clusters at zero or low grafting density (Figure 3a and Figure S1). Under the same conditions of high Mg^2+^ and low grafting density, the DLS data shows larger average *D*_h_ than under any other condition, again consistent with the formation of particle clusters. In the AFM images, it is possible to distinguish between individual particles and clusters. Quantifying the height of individual particles, values for *H* at low grafting density are very similar in the presence of Na^+^ or Mg^2+^. In contrast, quantifying the total particle volumes including clusters from AFM images, we find the largest average volumes for zero or low grafting density in the presence of Mg^2+^, with values much larger than for Na^+^ for the same grafting density (Figure 3h). Together, these results strongly suggest that the presence of Mg^2+^ tends to induce the formation of clusters and aggregation at low grafting density. In turn, high grafting of ssDNA increases the colloidal stability and protects the particles against aggregation, which is desirable for many applications.

### The effect of salt on the DNA shell

At high grafting density, both the Na^+^ and Mg^2+^ conditions show similar behavior. For a given high grafting density, we observe a consistent decrease in *D*_h_ going from low salt to 300 mM Na^+^and a further decrease in 5 mM Mg^2+^, in line with the expectation that a higher salt concentration leads to a tighter, less extended hydration layer and ion atmosphere.^43^ The magnitude of the changes of a few nm is similar to the change in Debye length (which is 3-4 nm for the low salt conditions and 0.6 nm for 300 mM NaCl) and the corresponding change in persistence length of ssDNA over this salt range.^45^ However, the fact that 5 mM Mg^2+^ leads to a larger reduction in *D*_h_ than 300 mM Na^+^, despite having a lower ionic strength, suggests that the divalent ions screen electrostatic interactions more efficiently than monovalent ions and lead to a tighter hydrodynamic shell.

Interestingly, for the highest grafting density, *H* and the particle volumes decrease in the presence of Mg^2+^, likely due to intra-strand ion bridging between ssDNA backbone charges and Mg^2+^ ions that induces compaction of ssDNA strands on the MNPs (Figure 3g). Similar behavior has recently been shown on Au NP-DNA superstructures.^46^ We derived a so-called compression-clustering propensity parameter by dividing *D*_h_ at high salt by *D*_h_ at low/no salt concentration (Figure 3i). In the presence of Mg^2+^, the particles reveal a sharp transition from shell compression to particle clustering by transitioning from the high to low DNA grafting density. In contrast, in the presence of Na^+^, *H* and the particle volumes increase further at the highest grafting density (Figure 3e and h), suggesting that the collapse observed in Mg^2+^ is specific to divalent ions and that Na^+^ appears to stabilize a dense DNA brush structure at high grafting density. The compression-clustering parameter is close to 1 in the case of Na^+^ for all grafting densities, indicating no salt-induced clustering or compression for the Na^+^ condition (Figure 3i).

### Intermediate grafting densities provide the highest sensitivity for magnetic biosensing

We next investigated whether the grafting density impacts the suitability of our CMP-DNA conjugates for applications, focusing on two promising assays: magnetic-based biosensing of nucleic acids^47^ and assembling on DNA origami. Our magnetic biosensing assay is based on monitoring changes in the magnetic relaxation dynamics of MNPs upon molecular interactions between the receptors on particles and target nucleic acids; it is simple, rapid, and offers a mix-and-measure sensing concept. We used magnetic particle spectroscopy (MPS)^48–50^ that monitors changes in the magnetic relaxation dynamics of MNPs in the higher harmonics spectrum.^51^ The MPS-based biosensing relies on the Brownian relaxation mechanism of MNPs, which depends on particle hydrodynamic volume. Upon functionalization of CMPs with DNA probes and hybridization with complementary sequences, the particle hydrodynamic volume increases, which in turn reduces the MPS harmonics amplitude, with the higher harmonics showing a larger relative reduction compared to the 3^rd^ harmonic. This indicates that the 5^th^ harmonic drops more than the 3^rd^ harmonic upon detecting the target sequence. Therefore, the ratio of the 5^th^ to 3^rd^ harmonics (HR53) of the particles drops upon sensing the targets compared to the HR53 of the particles without the targets.^52^ Our CMPs with a small *D*_h_ relaxing via Brownian processes are ideal for MPS-based magnetic biosensing, since small changes in the particle hydrodynamic volume are registered in the harmonics spectrum.^53^

For these assays, we labeled our CMP-DNA conjugates with a DNA sequence of 30 nucleotides (GC = 23.3%, *T*_m_ = 53 °C) at R(x) = 0.15, 0.48, and 1.2, corresponding to values below, at, and above the *R*_half_ value derived from the DLS data (Figure 1h; see Methods for details). To eliminate the effect of particle concentration on the assay results, we calculated the 5^th^ to 3^rd^ harmonics ratio (HR53) as a concentration-independent measure for the sensing capability (see Methods for details).^54^ We measure the change in HR53 prior (w/o) and after (w/) adding target complementary sequences and letting them base pair with the DNA strands on CMPs. All three samples show a drop in the HR53 ratio after hybridizing with the target sequences (Figure 4a). We find that the relative change in the HR53 upon sensing the target changes non-monotonously with DNA probe grafting density (Figure 4a). At both low and high ssDNA probe grafting densities, hybridization with the complementary sequences leads to a smaller relative change in the HR53 ratio compared to the intermediate grafting density, which shows the highest relative change in the HR53 upon sensing the target DNA (Figure 4b). We interpret the non-monotonous sensitivity to target DNA hybridization as follows: At low grafting density, there are only relatively few DNA molecules available for binding and, therefore, the effect of DNA hybridization on particle dynamics is small (Figure 4c). More surprisingly, the response to target DNA binding at the highest grafting density is also relatively small. The limited response at high grafting density might indicate that the grafted ssDNA forms dense polymer brushes on the CMPs at high density, such that the accessibility for the target DNA is limited or that a further change in particle hydrodynamic volume upon hybridization with the complementary sequences is limited. At intermediate grafting densities, near the midpoint of the Hill-like transition, the grafted DNA appears to form a mixture of coiled and brushed configuration on the particles. This configuration is readily accessible to the complementary sequences and upon hybridization induces a large change in particle hydrodynamic size and magnetic relaxation dynamics, providing optimal grafting density for sensing (Figure 4c).

**Figure 4.**
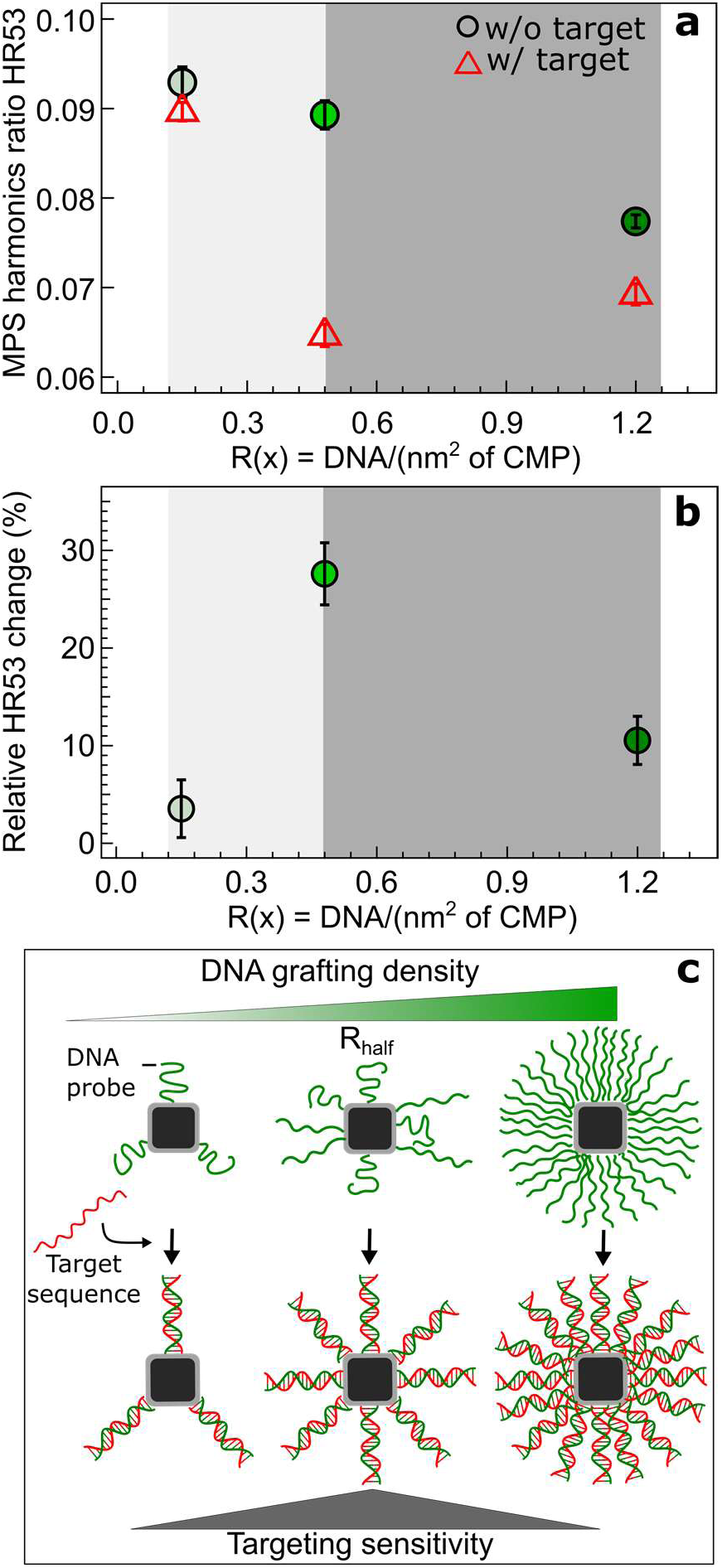
Magnetic particle spectroscopy-based biosensing of target nucleic acids. (a) MPS harmonics ratio HR53 without (w/o) and with (w/) complementary target DNA for three different grafting ratios. (b) Relative change in the HR53 upon sensing the target nucleic acids given by (HR53_probe_-HR53_probe+target_ / HR53_probe_), which is a measure of the assay sensitivity. The light and dark gray areas highlight the low and high grafting density regimes, respectively. (c) Schematic illustration of the organization of the DNA probes on CMPs depending on the grafting density. The MPS-based assays were performed at the DNA target-to-probe ratio of 5 at 500 mM NaCl concentration.

### High DNA grafting density favors efficient and site-specific coupling of magnetic nanoparticles to DNA origami structures

Next, we tested if the ssDNA grafting density is relevant for assembling CMP-DNAs on DNA origami nanostructures. For DNA origami binding experiments, we functionalized CMPs with 20-mer thymine (T_(20)_) strands and designed the DNA origami with 20-mer adenine (A_(20)_) as docking sites, since this combination is a well-established and robust protocol for patterning inorganic NPs on DNA origami. We had no success in coupling our CMP-DNA conjugates at low DNA grafting density to 6 helix bundle (HB) or 24 HB DNA origamis, mainly due to particle clustering. Our DNA origami binding experiments used 11 mM Mg^2+^ to stabilize the DNA origami, even higher than the Mg^2+^ concentration of 5 mM that was observed by DLS and AFM to promote clustering at low grafting density (Figure 3f). We, however, successfully assembled our CMP-DNA conjugates at R(x) = 1.2 with high efficiency on 6HB DNA origami that have ∼50 docking sites on three facets along the structure (Figure 5a). We find that our CMP-DNA conjugates assemble densely on the origami, with, on average, 12.5 CMPs per structure (Figure 5b), indicating a ∼20% binding efficiency. Of note that due to steric hindrance between neighboring particles on the closely positioned docking sites reaching 100% efficiency is nearly impossible. The CMPs with the highest DNA grafting density also enable site-specific binding to 24HB structures that have two binding sites at two ends of the structure (Figure 5c). The particle center-to-center distance determined from TEM images is 74.5 ± 4.6 nm (mean ± std), in excellent agreement with the distance of 80 nm expected from the origami design (Figure 5d). Our results demonstrate the importance of a dense ssDNA shell around MNPs for an effective assembly of MNPs on DNA origami nanostructures.

**Figure 5.**
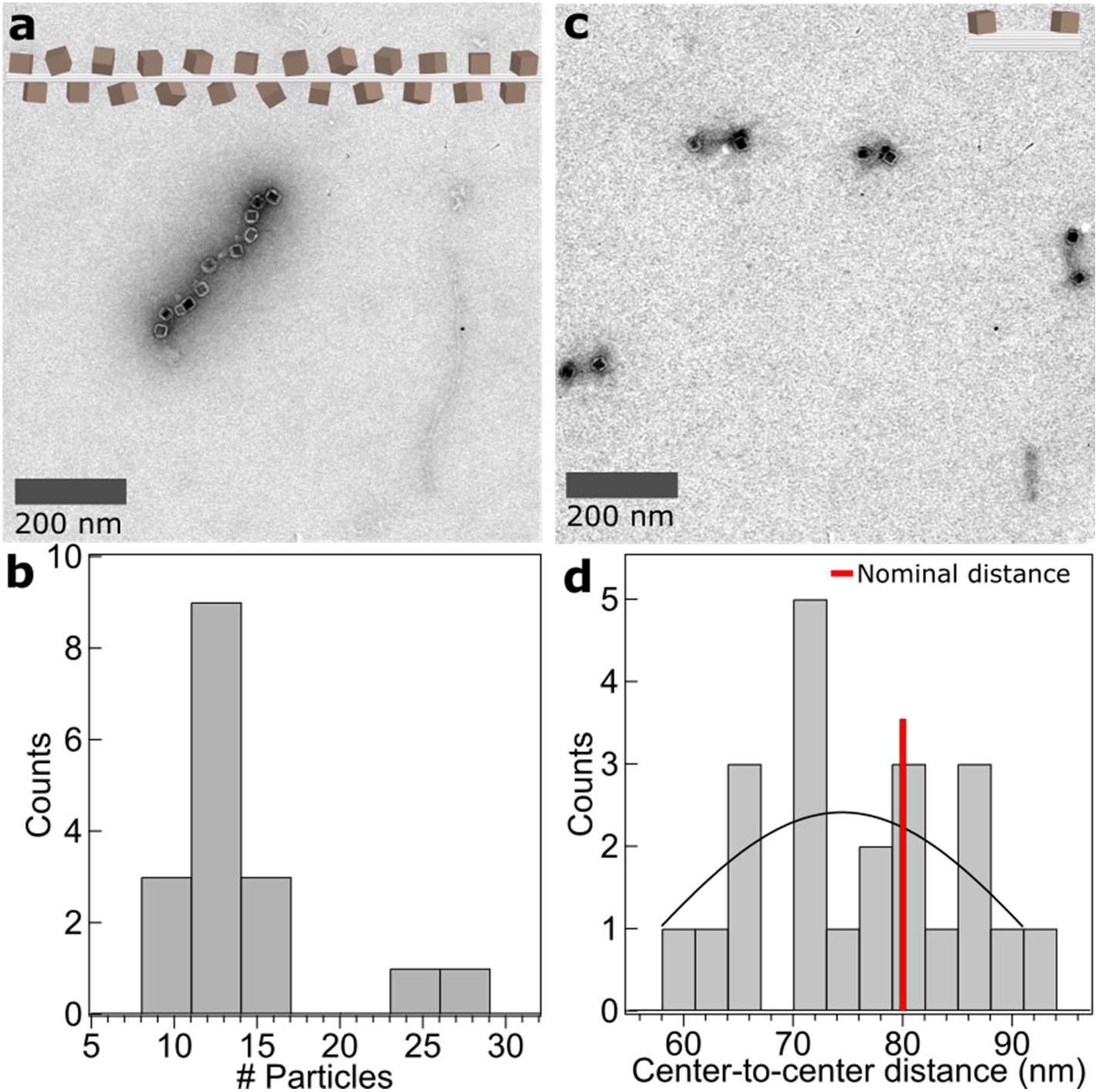
Negative-stain transmission electron microscopy analyses of the assembly of CMP-DNA conjugates on DNA origami nanostructures. (a) Assembly of CMPs labeled with T_(20)_ at a density of R(x) = 1.2 to 6HB DNA origami with ∼50 A_(20)_ docking sites on three facets along the structure. (b) Histogram of number of particles assembled on individual 6HB structures. (c) Site-specific binding of CMP-DNA(T)_20_ at R(x) = 1.2 to 24HB DNA origami with two A_20_ docking sites at two ends of the structure. (d) Histogram of particle center-to-center distance on the 24HB structures. The solid line is a Gaussian indicating a mean particle center-to-center distance of 74.5 ± 4.6 nm (mean ± std). The vertical red line is the distance between the docking sites expected from the DNA origami design.

## CONCLUSION

We show that our polymer encapsulated CMPs with high colloidal stability and small particle hydrodynamic size are a powerful platform to decipher effects of ssDNA grafting density on a range of dynamic properties of magnetic nanoparticles. We find that the magnetic relaxation, hydrodynamic, and electrophoretic mobilities change non-linearly with the number of ssDNA on CMPs, and can be well described with the Hill equation. The fits of Hill equations to gel electrophoresis mobility, dynamic light scattering, and magnetic ac-susceptibility data result in similar midpoint and rate parameters. At low grafting density, the ssDNA strands are mainly in a coiled configuration, in contrast to the situation at high grafting density, where the ssDNA form dense polymer brushes on CMPs. The brushed ssDNA strands on CMPs are compressed in the presence of divalent Mg^2+^ ions as observed in DLS and AFM analyses. We attribute this behavior to intra-DNA strand ion bridging, which is not seen in the presence of monovalent Na^+^ ions. Importantly, we find that intermediate to high DNA grafting densities enhance the colloidal stability of the CMPs and effectively prevent clustering and aggregation even in the presence of divalent ions.

Our results show that depending on the application of colloidal MNPs, different ssDNA grafting densities have to be aimed for. Using our CMP-DNA conjugates as magnetic nanomarkers for MPS-based magnetic biosensing of target nucleic acids, we find that the ssDNA probe grafting density at the midpoint of the Hill equation leads to the highest sensitivity of magnetic assays with MNPs. For coupling of CMPs to DNA origami structures, we demonstrate that the high grafting density of the ssDNA probe favors an efficient and site-specific coupling. Our polymer coating protocol with clickable polymers is a versatile approach to transfer organic ligand-coated MNPs in water. By applying our polymer coating and DNA labelling approaches to MNPs that can be detected at picomolar concentrations,^38^ we expect to achieve picomolar nucleic acid detection sensitivity. Our CMPs with a tunable DNA grafting density open up new opportunities, e.g. in combining magnetic biosensing with DNA switches such as DNA strand displacement and the realization of DNA origami structures with magnetic actuation capabilities.

## Methods

### Chemicals

Iron (III) acetylacetonate (99.9% trace metal basis), cobalt (II) acetylacetonate (≥99.0%), dibenzyl ether (DBE, ≥98%), oleic acid (OA, 90%), 1-octadecene (ODE, 90%), poly(maleic anhydride-alt-1-octadecene) (Mn=30.000-50.000), NH_2_-PEG(3)-Azide, Tris-Base, EDTA, MgCl_2_, NaCl were purchased from Sigma-Aldrich (Germany). All ssDNA oligonucleotides were obtained from Biomers GmbH (Germany). Sodium oleate (>97%) was purchased from TCI (USA). All organic solvents with ACS reagent-grading were purchased from Carl Roth (Germany) and used without further purification.

### Magnetic nanoparticle synthesis

Cobalt-doped iron oxide nanocubes were synthesized via thermal decomposition of iron (III) acetylacetonate and cobalt (II) acetylacetonate in a mixture of dibenzyl ether, 1-octadecene, oleic acid, and sodium oleate at 290 °C for 30 min as previously described.^38^

### Modification of poly(maleic anhydride-alt-1-octadecene) with NH_2_-PEG(3)-azide linker

Poly(maleic anhydride-alt-1-octadecene) (PMAO) was modified with NH_2_-PEG(3)-azide via anhydride ring opening reaction following the procedure published by Jin et al.^40^ with some modifications. In a typical reaction, 145 mg (0.482 mmol monomer units) of PMAO were added into 7.5 mL dimethylformamide (DMF) in a round flask and heated up to 65°C. Next, 100 mg (0.482 mmol) of NH_2_-PEG(3)-Azide dissolved in 1 mL DMF was pipetted into the reaction flask and left to react for 24 h at 65 °C. After cooling the flask to room temperature, DMF was removed using a rotary evaporator operating at 5 mbar and 36 °C. The crude yellowish oily product was dissolved in 3 mL of tetrahydrofuran (THF) and dialyzed against THF using regenerated cellulose dialysis tubes (6 kDa cutoff size) to remove unreacted PEG and byproducts. Next, THF was removed thoroughly in a rotary evaporator and the resulting product was dissolved in 9 ml of chloroform and stored at 4 °C prior to usage.

### Polymer coating procedure

The polymer coating of oleic acid coated NPs was performed following the protocol by Pellegrino et al.^55^ In a typical procedure, 5.7 ml (containing ∼ 35 mg/ml of PMAO-azide polymer) was poured into a round glass flask and sonicated for 10 min. Next, 2 ml of particle suspensions in chloroform (3 g(MNP)/l) were first further diluted with 5.5 ml of chloroform and then added dropwise into the polymer solution and sonicated again for 30 min to homogenize the mixture. The ratio of polymer to MNPs corresponds to 500 polymer units per nm^2^ of a particle. Afterwards, chloroform was removed very slowly within 5-6 h through stepwise reduction of pressure to a final value of ∼350 mbar and increase of temperature to 34 °C. After a complete removal of chloroform, particles were resuspended in 20 ml of sodium borate buffer (pH 8.7) by sonication for 1 h at ∼45 °C. The particle suspension was then concentrated to 2 ml using spin filtration (Amicon regenerated cellulose spin filter, 15 mL, 30 kDa cutoff size) at 3200 rpm at 20°C for 30 min. These particles offer click-chemistry conjugation and are named clickable magnetic nanoparticles (CMPs).

### Purification and particle fractionation

To remove polymeric micelles that are formed during the polymer coating process, we fractioned CMPs on non-continuous sucrose gradient (10%:40%:60%, from top to bottom, each fraction 4 ml) centrifuge columns. Typically, 500 µL of PMAO-coated particle suspensions in borate buffer were loaded into the sucrose centrifuge tube and centrifuged for 2 h at 4500 rpm at 4 °C. Next, singly polymer coated nanoparticles were collected in the 10% band using a long needle. The sucrose was then removed by 5 rounds of Amicon filtration (Amicon spin filter, 15 mL, 30 kDa cutoff size) at 3200 rpm and 20 °C for 12 min. After each round of spin filtration the particles were thoroughly resuspended in borate buffer by vigorous pipetting. Finally, the particles were concentrated to 2 mL of Tris-EDTA (5 mM Tris, 1 mM EDTA, 5 mM NaCl, pH 7.3) buffer by spin filtration.

To further purify the particles, we performed one round of magnetic washing. To do so, we used a 1.5 ml MACS column (Miltenyi Biotec) in combination with a MiniMACS separator (permanent magnet). First, the MACS column was placed in the separator and a collection vial was placed under the column to collect all effluent. At first, the column was rinsed with 500 μl of TE buffer and then the particle suspension was dropwise loaded on top of the column. After making sure that the whole buffer is passed through the column, the column was removed from the permanent magnet. Afterwards, the MNPs were collected in a glass vial by adding 3×565μl of TE buffer and pushing the plunger supplied with the column. The particle suspension was then stored at 4 °C prior to DNA functionalization. The particles after polymer coating are called clickable magnetic nanoparticles (CMPs).

### DNA labelling of CMPs

The following equation was used to calculate the amount of ssDNA (100 μM) needed to label CMPs with ssDNA strands of 20-mer thymine (T) at different R(x) = (ssDNA/nm^2^ of CMP) ratios:

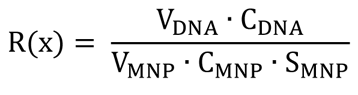

with *V*_DNA_ (µl)*, V*_MNP_ (μl), *C*_MNP_ (μM), and *C*_DNA_ (μM) the volume and concentration of the ssDNA solution and the CMP suspension, respectively. The surface area of a cubic particle is given by 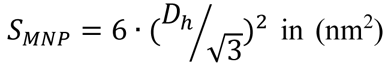, with *D*_h_ the volume weighted particle hydrodynamic diameter measured by DLS. Of note that the DLS analysis software assumes a spherical shape for particles.

In a typical experiment, a 180 µl of CMP suspension in TE buffer at a particle concentration of 112 nM was mixed with 112 µl, 224 µl, 447 µl l, 559 µl and 894 µl of 100 µM 5’-T(20)-DBCO-3’ ssDNA (100 μM in TE) to obtain R(x) of 0.15, 0.3, 0.6, 0.48, and 1.2, respectively. After mixing, the sample was left on bench at room temperature overnight. To maximize the binding efficiency of ssDNA to CMPs, we applied a so-called salt aging procedure using NaCl salt the next day.^56^ We increased the concentration of NaCl in the mixture incrementally by 100 mM every hour up to 400 mM. After each salt adjustment step, the mixture was homogenized by vortexing and sonicating for ∼20 s. At the end of the salt aging procedure, the mixture was left on bench at room temperature overnight. Next, the excess of ssDNA was washed out by 3 rounds of centrifugation at 14000 rpm, 5 °C, and for 13 min. Finally, the sample volume was adjusted again to 180 µl to obtain the original particle concentration. The samples were stored at 4 °C prior to further use.

### Magnetic particle spectroscopy (MPS)-based magnetic bioassays of target nucleic acids

We performed magnetic biosensing assays on a different sequence than 20-mer T to avoid unspecific cross-linking between CMPs upon sensing complementary sequences. For magnetic particle spectroscopy (MPS)-based magnetic biosensing assays, we labelled CMPs at R(x) = 0.15, 0.48, and 1.2 grafting ratios with 5′-ACT GCT TAT GCT AAT AGT GTA AAA AAA AAA-DBCO-3′ (30 nucleotides, GC = 23.3%). The sequence has 5A nt as a spacer.

The target sequence is: 5′-<u>TTTTTACACTATTAGCATAAGCAGT</u>TGTGGCATCTCCTGATGAGGTCTCTCTCTCTCT CTCTCTC-3′ The underlined section shows the complementary section to the DNA probe on CMPs. In these assays, the ratio of target DNA to ssDNA probe on CMPs was set to 5 to ensure enough excess of the target. Typically, the mixture of CMP-DNA conjugates and target DNA in TE buffer with 500 mM NaCl was shaken at 300 rpm for 2 h. Next, the mixture was kept at 4 °C overnight. The MPS measurements were then performed directly on the mixture in a wash-free fashion.

### DNA origami synthesis

DNA origami folding is performed in a one-pot reaction. First, all necessary ingredients were mixed together: the scaffold (p8064 for 24HB, p8634 for 6HB), staples and buffer (1x TE, 24mM MgCl_2_ for 24HB and 18mM MgCl_2_ for 6HB). Typically, we fold the scaffold with a 5x excess of the staples, so for a scaffold concentration of 20 nM, a staple concentration of 100 nM is used. The mixture was first heated up and then went through an annealing program (65 °C to 64 °C at 15min/°C, 64 °C to 59 °C at 5min/°C, 59 °C to 39 °C at 45 min/°C, 39 °C to 36 °C at 30 min/°C, and 36 °C to 20 °C at 5min/°C). The first purification of the origami was performed by PEG precipitation^57^ to increase the concentration before purifying it with an agarose gel (1%, 70 V, 80 min). The gel running buffer was 1xTAE with 11 mM MgCl_2_.

### Coupling of magnetic particles to origami

The CMP-DNA conjugates prepared at R(x) = 1.2 were mixed together with the purified DNA origami with an excess of 10:1 particles per binding site in 1xTAE (11 mM MgCl_2_) using the gel buffer conditions. The mixture went through the following temperature program: (40 °C to 20 °C at 10min/ °C, repeat 3 times). After gel purification, the sample was imaged with negative-stain TEM imaging.

### Characterization of magnetic nanoparticles

#### Transmission electron microscopy (TEM)

Transmission electron microscopy studies were carried out using a JEOL TEM microscope operating at 100 kV. The TEM samples were prepared by drop casting 5 µl particle suspensions in chloroform on a TEM grid (formvar-carbon coated copper grids with the mesh size of 300) and letting it completely dry in a fume hood. Negative-stain TEM samples were prepared by staining with uranyl formate (1%) for 10 s after 5 min incubation of particle suspension on TEM grid. The particle size *D*_c_ histogram was acquired by the analysis of a large field of view TEM image using the automatic particle analysis routine of the imageJ software.

#### Agarose gel electrophoresis shift assays

To study electrophoretic migration behavior of particles prior and after ssDNA labelling, native agarose gel electrophoresis (AGE) without staining was applied. Typically, 0.5% agarose gel in 1×TAE running buffer was made by dissolving 250 mg of agarose broad band gel in 50 ml of TAE buffer by 1 min microwave irradiation and 1 min magnetic stirring. The homogeneously dissolved gel solution was casted on a gel mold and left to solidify for 30 min. Afterwards, the casted gel was placed in electrophoresis chamber (Biorad) and samples were pipetted (containing 20% glycerol) into pockets. All AGE shift assays were run at 90 mA for 30 min. The gels were imaged under white and UV light exposure.

#### Analysis of gel images

To quantify the migration of particles on the gel, we used the Image Lab software of the Gel Doc instrument. The relative front *R*_f_ parameter was derived from these analyses, showing how far the particle band is moving in the gel relative to the length of the area that was selected as a band length. The *R*_f_ values were obtained using analysis tools available in the Image Lab software.

#### Depletion assays

To determine the number of ssDNA per particle at different nominal grafting ratios R(x), we measured the UV absorption at 260 nm on the supernatants after DNA functionalization and centrifugation of CMPs (14000 rpm, 4 °C, 13 min). To eliminate the background caused by ever existing CMPs in the supernatant, we set the absorption at 310 nm as the background. The concentration of ssDNA in the supernatant *c* in µg/l is calculated using the Beer-Lambert law given by:

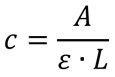

with *A* the absorption, *L* the optical path length in cm, *ε* the coefficient of excitation with *ε* = 0.027·10^-3^ (µg/l)^-1^·cm^-1^ for ssDNA. By plugging *c* into the following equation the average number of ssDNA per CMPs can be estimated.

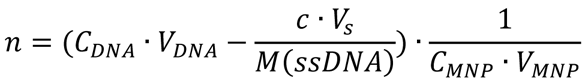

with *C*_DNA_ the DNA concentration in µM and *V*_DNA_ the volume of the DNA sample in µl, *V*_s_ the volume of the supernatant in µl*, M* the molecular weight of ssDNA (6250 g/mol for 20T), *C*_MNP_ the particle concentration in µM and *V*_MNP_ the volume of the particle suspension in µl. The results presented in Fig. 1h are the average value of three independent UV absorption measurements.

#### Dynamic light scattering (DLS)

DLS measurements were performed using a Malvern Zetasizer instrument at 173° backscattered measurement mode. Measurements were performed on particle suspensions in TE buffer (pH 7.3) at a typical particle concentration of ∼0.015-0.2 g/l at room temperature.

#### Sample preparation for atomic force microscopy (AFM)

For the AFM samples, we deposited 20 μl of CMPs or CMP-DNA conjugates at different grafting densities in TE buffer containing either 5 mM MgCl_2_ or 300 mM NaCl on freshly cleaved bare muscovite mica. The sample was incubated 5 min before washing with 20 ml MilliQ water and drying with a gentle stream of filtered argon gas.^58^

#### AFM imaging

The dry AFM images were recorded in tapping mode at room temperature using a Nanowizard Ultraspeed 2 (JPK, Berlin, Germany) AFM with silicon tips (FASTSCAN-A, drive frequency 1400 kHz, tip radius 5 nm, Bruker, Billerica, Massachusetts, USA). Images were scanned over different fields of view, with a pixel size of 1 nm/pixel and with a scanning speed of 5 Hz. The free amplitude was set to 20 nm. The amplitude setpoint was set to 80% of the free amplitude and adjusted to maintain a good image resolution.

#### AFM image analysis

Postprocessing of AFM data was performed in the software SPIP (v.6.4, Image Metrology, Hørsholm, Denmark) using the particle and pore analysis to determine the height of the nanoparticles. Contaminations (and for some parts of the analysis clusters) were excluded manually.

#### Complex ac-susceptibility (ACS) spectroscopy

The ACS spectra were recorded using our home-built AC spectrometer operating from 200 Hz to 1 MHz at magnetic field amplitudes of *µ_0_H* = 95 µT. The measurements were carried out at 295 K on 150 μl of particle suspensions at a particle concentration of 112 nM.

#### Magnetic particle spectroscopy (MPS)

The MPS measurements on particles in TE buffer were performed using a custom-built MPS setup (immunoMPS),^38^ which is especially designed for sensitive magnetic immunoassays. The setup operates at an excitation frequency of 590 Hz and a magnetic field amplitude of *µ_0_H* = 15 mT. The measurements were performed on 60 µl particle suspensions at a typical particle concentration of 20 nM and at a temperature of 295 K.

#### MPS-based magnetic bioassays and analysis of MPS harmonics ratio

Our magnetic assays are based on changes in Brownian magnetic relaxation processes of MNPs that happen when the particle hydrodynamic size/volume increases upon hybridization of the DNA probes on the particles with the target DNA sequences in solution. A molecular binding event slows down the Brownian relaxation of the particles in ac magnetic fields and thereby reduces the amplitude of the MPS harmonics. The MPS higher harmonics are, however, more sensitive to changes in the particle relaxation dynamics than the fundamental excitation frequency. This means that the 5^th^ harmonic drops more than the 3^rd^ harmonic upon the binding event. Therefore, the ratio of the 5^th^ to 3^rd^ harmonics (HR53) drops after a successful binding to the target sequence compared to the HR53 of the particle sample without the target sequence. The MPS harmonics ratio is, therefore, a particle concentration-independent measure to judge success of MPS-based assays.

#### Cooperative dynamics: Hill equation

Relative gel front *R*_f_, hydrodynamic size *D*_h_, and the ACS peak frequency *f*_p_ vs. R(x) were fitted with a Hill equation given by:

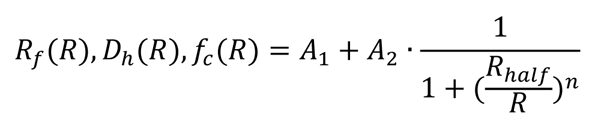

*A_1_, A_2_*, *n*, and *R*_half_ are fitting parameters. *R*_half_ is the nominal grafting density that produces a half-maximal change in the respective parameters and *n* is the Hill coefficient. For the fit to the *R*_f_ data the parameter *A_1_* is set to zero.

#### Data Availability

Uncropped agarose gel, TEM, and negative-stain TEM images and source data for AFM and DLS data are available.

#### Supporting Information

Figure S1, S2, and S3 are given.

#### Corresponding Author

Aidin Lak

#### Author Contributions

A.L. conceptualized and designed the research as Alexander von Humboldt fellow at LMU Munich, synthesized the polymer, established polymer coating and DNA functionalization protocols, performed TEM and DLS measurements and analyses, modelled the ACS data, supervised the study, analyzed the data, and wrote the manuscript. Y.W. performed polymer coating, DNA functionalization, and hybridization assays, AGE, ACS, MPS, and DLS measurements. P.J.K. prepared AFM samples, performed measurements, and analyzed data. C.P. prepared the DNA origami, coupled particles to DNA origami, performed negative-stain TEM imaging, and analyzed the data. M.S.C. synthesized the particles. M.C. co-developed the polymer coating procedure. F.L. and T.V. developed ACS and MPS setups. F.S. P.T., M.S., T.L., J.T. analyzed the data. J.L. analyzed the data, supervised the study at LMU Munich, and wrote the manuscript. All authors have given approval to the final version of the manuscript.

## Supporting information

Supporting Informtion

## Acknowledgment

This work is supported by DFG RTG 1952 “NanoMet”, DFG LA 4923/3-1, Junior Research Group “Metrology4life”, and the Alexander von Humboldt Foundation. J.T. is funded under DFG TA 1375/1-1 and C.P. and T.L. through the DFG SFB1032 “Nanoagents,” Project A6. We thank Margherita Gallano and Gloria Müller for the initial characterization of DNA labelled MNPs. We thank Kerstin Frank, Petra Schmidt (TU Braunschweig) for ICP-OES measurements, Willem Vanderlinden for helpful discussions about AFM imaging, and Thomas Nicolaus for laboratory assistance.

