## Supporting Informtion for "Cooperative dynamics of DNA grafted magnetic nanoparticles optimize magnetic biosensing and coupling to DNA origami"

**Content:**

**Supplementary Figure S1**

**Supplementary Figure S2**

**Supplementary Figure S3**

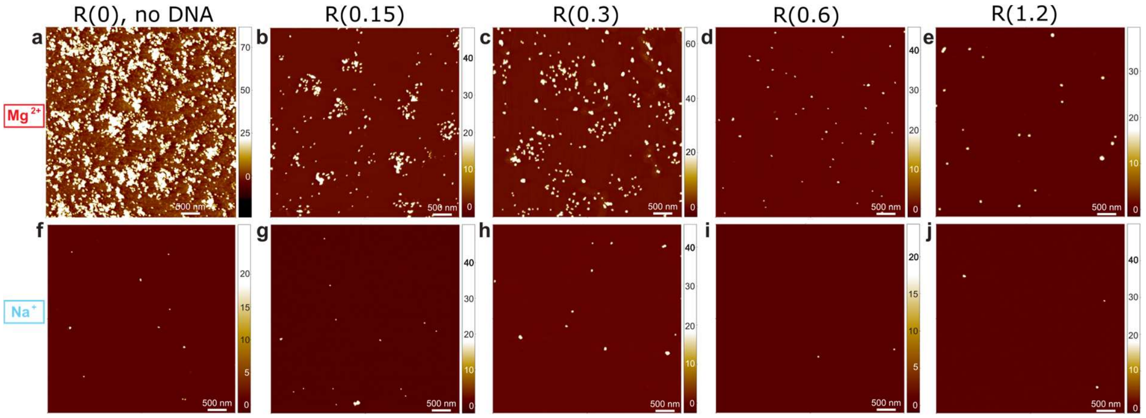


**Figure S1.** Atomic force microscopy height images of CMPs with no ssDNA grafting and of CMP-DNA conjugates at different grafting densities in the presence of 5 mM $MgCl_2$ divalent salt (top row) or 300 mM NaCl monovalent (bottom row). AFM images were recorded in tapping mode using a FASTSCAN-A cantilever on bare mica after drying. The z-ranges are indicated by the color map on the right side of each image (in nm).

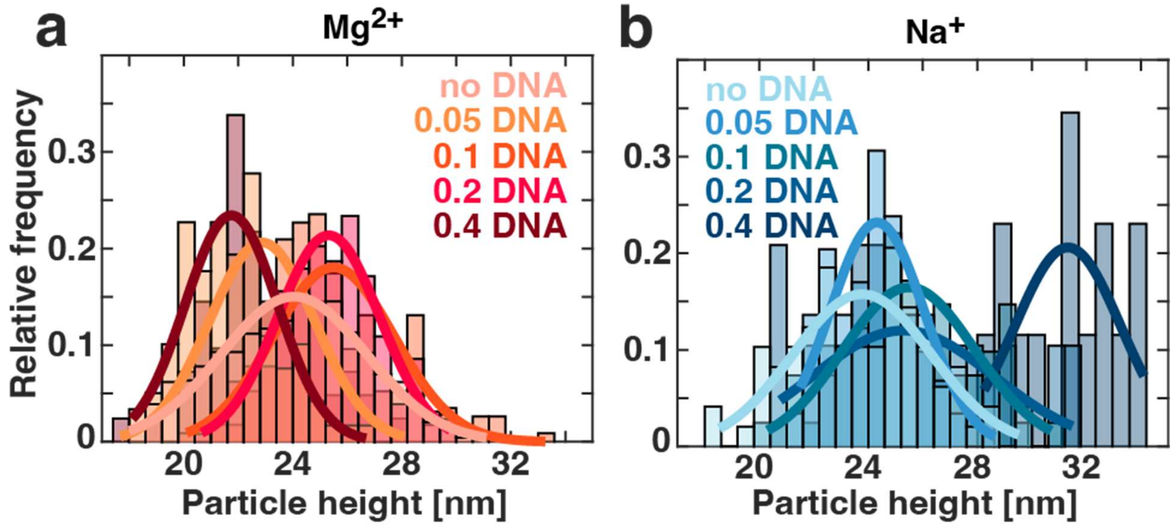


**Figure S2.** Normalized particle height histograms of CMPs with no DNA grafting and of CMP-DNA conjugates at different grafting densities indicated in the legend. Data are in the presence of (a) 5 mM $MgCl_2$ and (b) 300 mM NaCl. The solid lines are Gaussian fits. Examples of the corresponding AFM images are shown in Figure S1. The mean ± standard deviation of the height distributions are shown in Figure 3e in the main text.

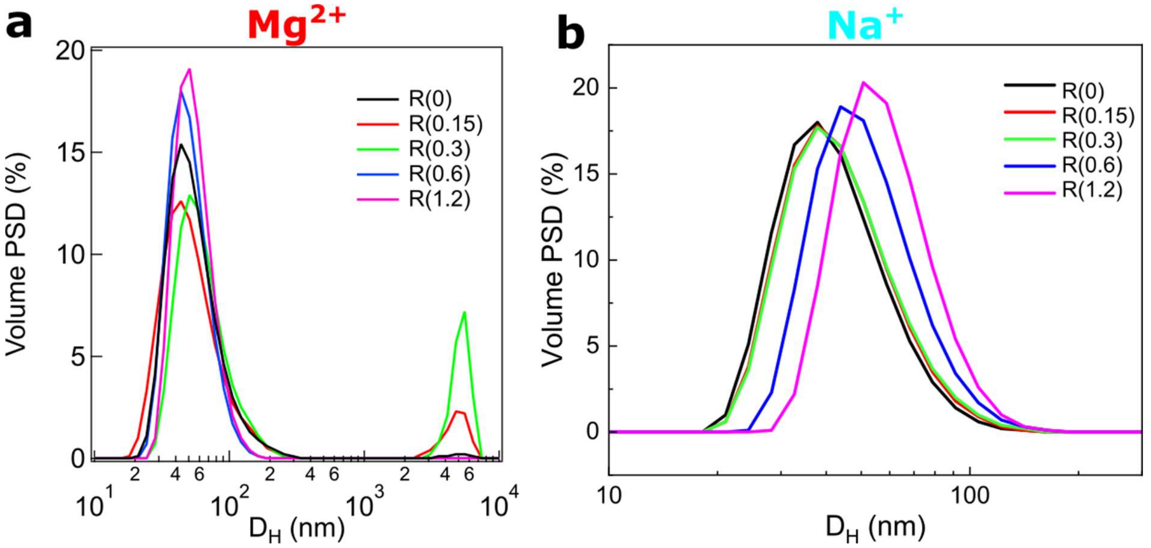


**Figure S3.** Volume-weighted particle size distribution (PSD) of CMPs with no DNA grafting R(0) and of CMP-DNA conjugates at all grafting densities measured by DLS in the presence of (a) 5 mM $MgCl_2$ divalent ions and (b) 300 mM NaCl monovalent ions.
